# Rapidly characterizing the fast dynamics of RNA genetic circuitry with cell-free transcription-translation (TX-TL) systems

**DOI:** 10.1101/003335

**Authors:** Melissa K. Takahashi, James Chappell, Clarmyra A. Hayes, Zachary Z. Sun, Vipul Singhal, Kevin J. Spring, Shaima Al-Khabouri, Christopher P. Fall, Vincent Noireaux, Richard M. Murray, Julius B. Lucks

**Author notes:** Corresponding Author: Julius B. Lucks.

## Abstract

RNA regulators are emerging as powerful tools to engineer synthetic genetic networks or rewire existing ones. A potential strength of RNA networks is that they may be able to propagate signals on timescales that are set by the fast degradation rates of RNAs. However, a current bottleneck to verifying this potential is the slow design-build-test cycle of evaluating these networks *in vivo*. Here we adapt an *Escherichia coli*-based cell-free transcription-translation (TX-TL) system for rapidly prototyping RNA networks. We used this system to measure the response time of an RNA transcription cascade to be approximately five minutes per step of the cascade. We also show that this response time can be adjusted with temperature and regulator threshold response tuning. Finally we use TX-TL to prototype a new RNA network, an RNA single input module, and show that this network temporally stages the expression of two genes *in vivo*.

## Introduction

A central goal of synthetic biology is to control cellular behavior in a predictable manner ^1^. Natural cellular behavior is governed by the expression of specific sets of genes needed for survival in different environments or in developmental life stages. Genetic networks – webs of interactions between cellular regulatory molecules – are responsible for dynamically turning these genes on at the right time and place, and in effect are the circuitry that implement behavioral programs in cells ^2^. Because of this, a central focus of synthetic biology has been to control cellular behavior by engineering genetic networks from the bottom up ^1^.

Historically, work on engineering genetic networks has focused on combining sets of regulatory proteins to control their own expression in patterns that implement behaviors such as bistable memory storage ^3^, oscillations ^4–6^, layered logic gates ^7,8^, advanced signal processing ^9^, and spatial control of gene expression ^10^. More recently, non-coding RNAs (ncRNAs) have emerged as powerful components for engineering genetic networks ^11^. There are now examples of engineered ncRNAs that regulate nearly all aspects of gene expression ^11–18^, some as a function of intracellular molecular signals^14,15,19^. In addition, new RNA structural characterization tools are enabling the engineering and optimization of these mechanisms ^11,19–22^. There are even large libraries of orthogonal RNA regulators ^21–23^, and there have been initial successes in engineering small genetic networks out of RNA regulators ^16,18,24,25^.

RNA genetic networks have several potential advantages over their protein counterparts 11. First, networks constructed from RNA-based transcriptional regulators propagate signals directly as RNAs, thus eliminating intermediate proteins and making them potentially simpler to design and implement ^18^. Second, tools based on qPCR and next-generation sequencing have the potential to characterize the species, structural states, and interactions of RNAs across the cell at a level of depth and detail not possible for proteins ^11^. Finally, since the speed of signal propagation in a network is governed by the decay rate of the signal ^26^, RNA networks have the potential to operate on timescales much faster than proteins. However, the design principles for engineering RNA circuitry are still in their infancy, and we have yet to fully test and verify these potential benefits. This is in part due to the slow nature of the current design-build-test cycle for engineering genetic networks *in vivo* that takes on the order of days even when testing circuits in fast growing bacteria.

Recently, cell-free protein synthesis systems have been developed into a platform to rapidly characterize the outputs of genetic networks ^27–33^. Cell-free reactions often consist of three components: cell extract or purified gene expression machinery, a buffer/energy mix optimized for gene expression, and DNA that encodes the genetic network ^27,34^ (Figure 1). Fluorescent proteins are generally used as reporters, thus monitoring fluorescence over time allows the characterization of circuit dynamics. Because of their simplicity, cell-free reactions reduce the time for testing a constructed genetic circuit design from days to as little as an hour ^32,35^. Since these systems do not require selection markers or DNA replication to maintain circuitry constructs, there are no limitations on DNA circularization or on plasmid origin of replication and antibiotic compatibility ^32,35^. This flexibility allows for faster, multiplexed generation of circuit constructs, further reducing design-build-test cycle times. Since cell-free reactions lack a membrane, DNA encoding different regulators can be added at any time during the reactions, enabling the rapid characterization of network response as a function of perturbations that are extremely difficult or even impossible to do inside cells ^33^ (Figure 1). Additionally, there is increasing evidence that these *in vitro* characterizations correlate to *in vivo* results, including comparable rates of RNA degradation ^28,32,35^. Cell-free systems thus have intriguing potential to serve as an intermediate layer to rapidly prototype circuit design and response before porting the designs to the more complex environment of the cell.

**Figure 1.**
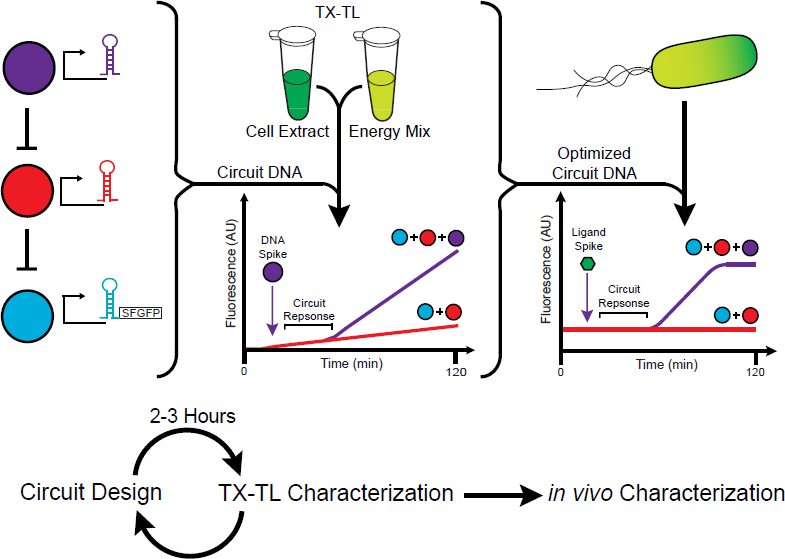
Schematic of the TX-TL design-build-test cycle for RNA circuits. Potential circuit designs are rapidly characterized in TX-TL by combing DNA-encoding RNA circuit components (colored circles) with the TX-TL reaction components. Overall circuit performance is monitored via the expression of fluorescent proteins enabling circuit designs to be rapidly benchmarked within a 2-3 hour period. In addition the openness of the TX-TL system allows characterization of circuit response via the addition of DNA encoding RNA regulators. After multiple iterations of the design-build-test cycle, optimized circuit designs can be transformed into E. coli and tested for in vivo functionality

In this work, we adapt an *E. coli* cell-free transcription-translation (TX-TL) system ^27,36^ for characterizing RNA genetic networks. Since this system was initially developed and optimized to test protein-based circuits ^28^, we start by validating the functionality of RNA transcriptional attenuators ^18^ in TX-TL, and characterize the effect of different TX-TL experimental conditions including DNA concentration and batch-to-batch variation. We then show that a double-repressive RNA transcriptional cascade functions in TX-TL with characteristics that match its *in vivo* performance ^18^. The ability to spike in DNA encoding the top level of this cascade during the reaction allowed us to directly probe the response time of this RNA network. We found that the response time of this RNA cascade is ∼5 minutes per step of the cascade, matching our expectation of quick signal propagation due to the fast kinetics of RNA degradation. We then show that this response time can be tuned by either changing the temperature, or effectively changing the threshold required for transcriptional repression by using tandem attenuators ^18^. To create a bridge to circuitry that we can implement and test *in vivo*, we show that we can use TX-TL to characterize the response time of similar cascades that use RNA regulators responsive to theophylline ^19^ to activate the cascade. The success of these experiments led to the forward design of a new RNA network motif, the single input module (SIM), which is responsible for staging the successive expression of multiple genes in natural pathways ^37^. After characterizing the functionality of the individual SIM components in TX-TL, we transfer the final RNA-SIM circuit to *E. coli*, and show that this network dynamically stages the expression of two fluorescent reporter proteins *in vivo*, solidifying the use of TX-TL for engineering RNA genetic circuits.

## Results and Discussion

### An RNA transcriptional cascade functions in TX-TL

In order to assess the feasibility of using TX-TL for characterizing RNA circuitry, we first tested the basic functionality of the central regulator in our RNA cascade, the pT181 transcriptional attenuator ^18^. The pT181 attenuator lies in the 5’-untranslated region of a transcript, and functions like a transcriptional switch by either allowing (ON) or blocking (OFF) elongation of RNA polymerase ^38,39^. The OFF state is induced through an interaction with an antisense RNA, expressed separately in our synthetic context ^18^ (Supporting Figure S1). By transcriptionally fusing the pT181 attenuator to the super folder green fluorescent protein (SFGFP) coding sequence, we are able to assess functionality of the attenuator by measuring SFGFP fluorescence with and without antisense RNA present (Supporting Figure S1).

To characterize antisense-mediated transcriptional repression in TX-TL, we first titrated concentrations of the pT181 attenuator reporter plasmid to determine an appropriate level of SFGFP output for our experimental setup. As expected, we observed a greater fluorescence output with increasing attenuator reporter plasmid concentration (Supporting Figure S2A). As noted in previous work with TX-TL, excessive amounts of input DNA can lead to resource competition and resource limitation within the reaction, which can confound circuit characterization ^27,29,33,36^. We found that an attenuator plasmid concentration of 0.5 nM (which corresponds to about one copy of plasmid into one *E. coli* cell) struck a balance between fluorescence signal and DNA concentration, and this concentration was used in subsequent experiments.

To test basic repression of the attenuator, we then characterized reactions that contained 0.5 nM of the attenuator reporter plasmid, and either 8 nM of the antisense-expressing plasmid (+), or 8nM of a control plasmid that lacked the antisense coding sequence (-) (Supporting Figure S3, Supporting Table S3-4). As expected, we observed a significant difference in the fluorescence trajectories between the (+) and (-) antisense conditions, with the (+) antisense condition resulting in an overall lower fluorescence output over time (Figure 2A). We note that in these experiments, we never observed a constant steady-state fluorescence signal due to the fact that SFGFP is not degraded (or is not diluted) in the TX-TL reaction during the timescale of the experiment – i.e. because SFGFP is not degraded, we always observed an increase in fluorescence over time even in the (+) antisense repressive condition.

**Figure 2.**
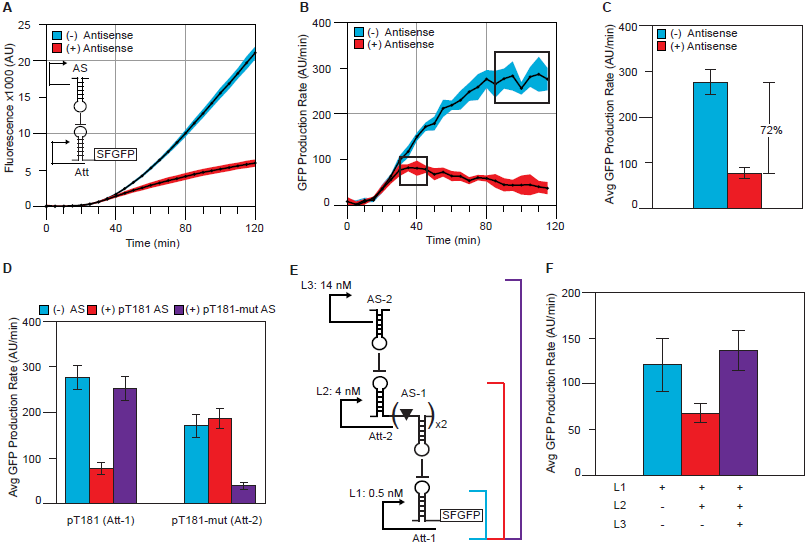
Characterizing RNA transcriptional attenuators and circuits in TX-TL. (A) Fluorescence time courses of TX-TL reactions containing the pT181 attenuator reporter plasmid at 0.5 nM, with 8 nM antisense plasmid (+) or 8 nM control plasmid (-). (B) SFGFP production rates were calculated from the data in (A) by calculating the slope between consecutive time points. Boxes represent regions of constant SFGFP production. Blue and red shaded regions in (A) and (B) represent standard deviations of at least seven independent reactions performed over multiple days calculated at each time point. (C) Average SFGFP production rates were calculated from the data in boxed regions in (B). Error bars represent standard deviations of those averages. The (+) antisense condition shows 72% attenuation compared to the (-) antisense condition in TX-TL. (D) Orthogonality of the pT181 attenuator (Att-1) to a pT181 mutant attenuator (Att-2). Average SFGFP production rates were calculated as in (C). Plots of SFGFP production rates can be found in Supporting Figure S2. Bars represent each attenuator at 0.5 nM with 8 nM of control plasmid (blue), pT181 antisense plasmid (red), or pT181 mutant antisense plasmid (purple). (E) Schematic of an RNA transcriptional cascade. L1 is the same pT181 attenuator (Att-1) reporter plasmid used in (A) – (D). In the plasmid for L2, the pT181-mut attenuator (Att-2) regulates two copies of the pT181 antisense (AS-1), each separated by a ribozyme (triangle) 18. The L3 plasmid transcribes the pT181-mut antisense (AS-2). (F) Average SFGFP production rates for the three combinations of the transcription cascade depicted (E). L1 alone (blue bar) leads to high SFGFP production. L1+L2 (red bar) results in AS-1 repressing Att-1, thus lower SFGFP production. L1+L2+L3 (purple bar) results in a double inversion leading to high SFGFP production. Total DNA concentration in each reaction was held constant at 18.5 nM. In (D) and (F) error bars represent standard deviations from at least seven independent reactions performed over multiple days.

Because of resource depletion effects that accumulate over time in TX-TL reactions ^27^, especially after 1-2 hours of incubation, endpoints of these fluorescence trajectories can give misleading quantifications of repression. Since the action of the antisense RNA ultimately affects the rate of transcription of the attenuator, we sought a way to quantify antisense repression by comparing the rates of SFGFP expression. Plotting the time derivative of the fluorescence trajectories allowed us to directly quantify the effective rate of SFGFP expression in the (+) and (-) antisense conditions as a function of time (Figure 2B). To further quantify antisense-mediated repression from these experiments, we computed a time average of the regions of constant maximum SFGFP production rate for each condition (Figure 2B boxed regions). We used maximum rate to reduce the confounding effects of resource depletion, which can cause production rates to go down over time (Figure 2B). Using these rates, we determined that with 8 nM of antisense RNA plasmid, we achieve 72% attenuation (Figure 2C), which is comparable to the 85% steady-state repression level previously observed *in vivo* ^18,23^.

In order to move toward characterizing RNA circuitry in TX-TL, we needed to confirm the functionality of an orthogonal attenuator/antisense pair. We used a mutant pT181 attenuator that had previously been shown to be perfectly orthogonal to the wild type attenuator *in vivo* ^18^. To test orthogonality in TX-TL, we performed reactions containing one of the two attenuator reporter plasmids at 0.5 nM, and one of the two antisense plasmids or the control plasmid at 8 nM. For both attenuators, we found results consistent with those from *in vivo* experiments ^18^ – cognate antisense RNAs caused repression, while non-cognate antisense yielded SFGFP expression rates that were within the error of the no-antisense conditions (Figure 2D, rate plots Supporting Figure S2 B-D). We thus confirmed the orthogonality of the two attenuator/antisense pairs in TX-TL.

Interestingly, we observed that the region of maximum SFGFP production occurs at different times for different combinations of antisense-attenuators in the reaction. In particular, reactions with cognate (repressive) antisense-attenuator pairs have maximum that occur around 40 minutes, while reactions with non-cognate (orthogonal), or just attenuator-reporter plasmids, have regions that occur near the end of the reactions at ∼ 100 min (Figure 2B, Supporting Figure S2C). Furthermore, cognate pairs show a decrease in reaction rate after the maximum is reached. The antisense-attenuator interaction is hypothesized to generate a double-stranded RNA complex ^40^, which is typically degraded by ribonuclease III (RNAse III). RNAse III itself is known to require magnesium ^41^, which is a key component for protein production in TX-TL ^36^. Thus the formation of the antisense-attenuator complex could end up increasing the rate of resource depletion in TX-TL reactions or accelerating the accumulation of by-products of the reaction, which inhibits TX-TL as well. This further motivates our use of the maximum SFGFP expression rate when quantifying repression in TX-TL.

The orthogonality of the two antisense-attenuator pairs allowed us to characterize a double-repression RNA transcriptional cascade in TX-TL (Figure 2E) ^18^. The bottom level of the cascade (L1) consists of an SFGFP coding sequence controlled by the pT181 attenuator’s (Att-1) interaction with its antisense (AS-1). AS-1 transcription is in turn controlled by the mutant antisense (AS-2), via a mutant attenuator sequence (Att-2) present upstream of AS-1 on the middle level of the cascade (L2). AS-2 is transcribed from the top level of the cascade (L3) (Figure 2E).

Previous work encoded L2 and L3 of the cascade on a high-copy plasmid (ColE1 origin, ∼200 copies/cell), and L1 on a medium copy plasmid (p15A origin, ∼15 copies/cell), and showed that the double repression cascade yielded a net activation of SFGFP expression at steady-state *in vivo* ^18^. Since there are no plasmid incompatibilities in TX-TL, we were able to use three separate plasmids for L1, L2, and L3 to characterize cascade function. Since three DNA elements are needed for the cascade, we first performed titrations of L2 versus 0.5 nM of L1 in order to find conditions that allowed us to observe repression without severe resource depletion (Supporting Figure S2E). We found that 4 nM of L2 caused a 72% repression, with no greater repression observed with higher concentrations of L2. To test the full cascade, we titrated different amounts of L3, from 10 nM to 18 nM, keeping the total amount of DNA in the TX-TL reaction constant with the addition of a control plasmid to approximately control for resource usage across conditions (Supporting Figure S2F). We found that L3 does in fact activate SFGFP expression with a rate that matches that of just L1 (Figure 2F).

This result proved that RNA circuitry functions in TX-TL reactions. Stated another way, these experiments demonstrate that the RNA circuitry tested only requires the machinery from the cytoplasmic extract contained in the TX-TL reaction. Furthermore, the flexibility of TX-TL allowed us to systematically validate the functioning of each level of the cascade by adding successive levels one at a time. This is in contrast to the previous *in vivo* work where complex controls were needed to validate cascade performance since L2 and L3 were encoded on the same plasmid ^18^. Finally, we note that all of these experiments were performed at 29°C, confirming that RNA transcriptional regulation and circuitry can function over a range of temperatures.

## Ideal TX-TL batch characteristics for circuit testing

There is known to be batch-to-batch variation in TX-TL reactions ^27^. In order to assess the impact of batch-to-batch variation on RNA circuit characterization, we tested three different extract/buffer preparations by adding a range of antisense control to 0.5 nM of the L1 plasmid. Since extra control DNA causes resource competition, this experimental design allowed us to assess the maximum amount of DNA per reaction that each batch could support. As shown in Figure 3, we observed several important features. First, for a fixed concentration of L1 and control DNA, we observed significant differences in the fluorescence time courses between the batches (Figure 3A). The endpoint fluorescence of batch 2 is more than twice that of batch 1 and 3. Second, batch 2 reaches constant SFGFP production faster than batch 1 and 3 for all conditions tested (Supporting Figure S4). Third, batch 1 had a lower fluorescence output than batch 3, but both reached constant SFGFP production at approximately the same time over all conditions.

**Figure 3.**
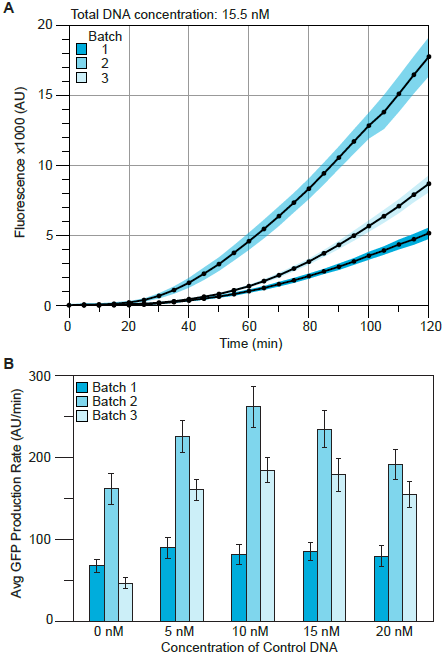
Assessing batch-to-batch variation. (A) Fluoresscence time courses of TX-TL reactions in three different extract and buffer preparations with 0.5 nM L1 and 15 nM control DNA. Shaded regions represent standard deviations of at least 11 independent reactions over multiple days calculated at each time point. (B) Average maximum SFGFP production rates for the same three buffer and extract preparations from reactions with 0.5 nM L1 and 0, 5, 10, 15, and 20 nM control DNA. Plots of maximum SFGFP production rates from which these were calculated can be found in Supporting Figure S4. Error bars represent standard deviations from at least 11 independent reactions.

In terms of DNA loading effect, in all of the batches we see an increase in production rate when adding 5 nM of control plasmid (Figure 3B). For batches 1 and 3, production rates remained constant up to the maximum concentration of control DNA tested (20 nM). The production rate for batch 2 increased with the addition of 10 nM control DNA and then decreased for 15 and 20 nM control DNA (Figure 3B).

We hypothesize that the increase in production rate for all batches upon adding 5 nM control DNA is due to RNA degradation machinery (RNAses) competition effects. The control DNA has the same promoter as the attenuator and antisense plasmids from which an RNA terminator is made (Supporting Figure S3). While this RNA does not affect the attenuator in a mechanistic way, it does provide a decoy for RNAses, and could cause a decrease in degradation rate of the attenuator-SFGFP mRNA. The increase in production rate from 5 to 10 nM control DNA in batch 2 could be the result of a higher RNAse content in that batch, thus more DNA is required to fully occupy the RNAses. Because of this loading effect, we found it important to use an appropriately constructed control DNA when designing comparative experiments (Figure S3).

Additionally, these results show the importance of screening TX-TL extract and buffer batches to best match the needs for circuitry characterization. For our RNA attenuator circuitry, we chose a batch that struck a balance between high production rate for better signal/noise, and invariance to DNA loading effects for testing circuitry with multiple components. We therefore used batch 3 in all further experiments.

### Characterizing the dynamics of RNA circuitry with TX-TL

One of the potential advantages of RNA over protein circuitry is a faster response time due to the relatively fast degradation of RNA molecules ^11^. The flexibility of TX-TL allowed us to directly measure the response time of the RNA transcription cascade (Figure 4). Since TX-TL is an open system, we designed an experiment that involved spiking in the DNA encoding L3 of the cascade into an ongoing reaction that was already expressing L1 and L2. We define the response time of this circuit, τ, as the time it takes to turn ON SFGFP production after spiking in L3. In order to determine τ, TX-TL reactions were setup with 0.5 nM L1 and 4 nM L2 following our earlier results (Figure 2). This reaction was allowed to proceed for 25 minutes, at which time (t=0) we spiked in 14 nM of either L3, or our control plasmid. Fluorescence trajectories showed that the L3 spike caused a noticeable deviation from the control trajectory ∼20 minutes after the spike (Figure 4B). By using Welch’s t-test to find the point at which the two trajectories differed significantly (Supporting Figure S5, Methods), we were able to quantify the response time over multiple experimental replicates to be 18.2 ± 6.0 min (Figure 4B).

**Figure 4.**
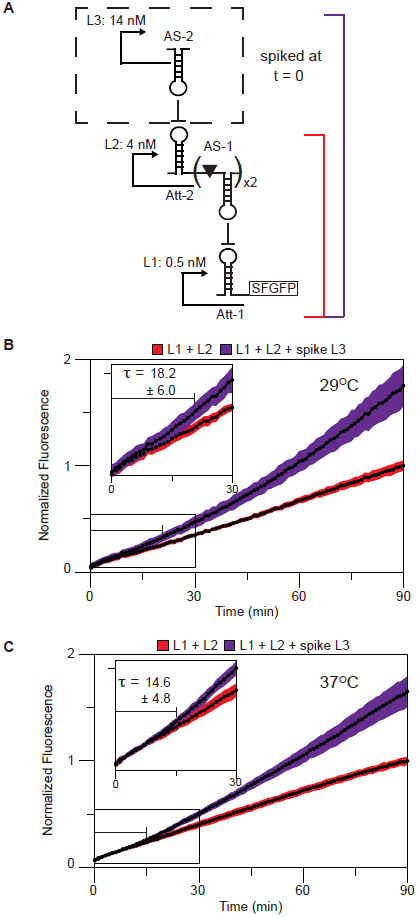
Determining cascade response time. (A) Schematic of spike experiment. L3 (or the no antisense control plasmid) was spiked into an ongoing L1+L2 TX-TL reaction at time, t = 0 (represented by dashed box). Concentrations of DNA used are indicated beside the levels. (B) Normalized fluorescence curves combining three separate experiments performed at 29°C with a total of 8 replicates over multiple days. L1 (0.5 nM) + L2 (4nM) reaction was setup for 25 min at which point L3 (14 nM, puple curve) or control DNA (14 nM, red curve) was spiked into the reaction and time reset to 0. Inset shows the response time of the circuit to the addition of L3; defined as the time at which the L3 spike curve is statistically different from the L1+L2 curve (τ = 18.2 ± 6.0 min). (C) Normalized fluorescence curves combining three separate experiments performed at 37°C with a total of 11 replicates over multiple days. The same experiment was setup as in (B) except that the L1+L2 reaction ran for 20 min prior to the addition of L3. τ = 14.6 ± 4.8 min. Shaded regions represent standard deviations calculated at each time point.

We can estimate τ by considering the three molecular events that need to occur to turn ON SFGFP production: i) AS-2 needs to be transcribed from L3, ii) the concentration of AS-2 needs to build up in order to repress the transcription of AS-1 via an interaction with Att-2, and iii) any existing AS-1 must be degraded. Using a simple ordinary differential equation model of the expression of each level of the RNA cascade, we can derive an expression for the response time (Supporting Appendix 1):

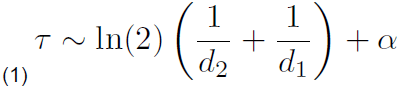

Where d_2_ and d_1_ are the degradation rates of AS-2 and AS-1, respectively, and α is the maturation time of SFGFP. Since ln(2)/d is the half-life of each antisense species, we find that the response time is a sum of half-lives of the intermediate RNA signals, similar to a related analysis of response times of protein circuitry ^2,26^. Using an estimate of a 5-minute half-life for our antisense RNAs ^42^ and a 5 minute maturation time for SFGFP ^43^, we can estimate τ from Equation (1) to be 15 minutes, in close agreement with our experimental observation.

Since antisense degradation rates set the timescale of Equation 1, we expect its rate to increase with increasing temperature, and thus expect increasing temperature to decrease τ. To test this, we repeated the same DNA spike experiment with reactions running at 37°C, and determined τ to be 14.6 ± 4.8 min (Figure 4C). This is in remarkable agreement to our 15 minute estimate of τ from Equation 1, which was derived from antisense degradation rates measured at 37°C. While the average response time at 37°C was lower than that at 29°C, the difference between the two averages was not statistically significant due to the large error bars on the measurements (p = 0.1694, Supporting Table S2). We believe with further measurements a statistically significant difference would be observed.

These results represent the first measurement of RNA circuitry response times. Furthermore, they confirm our expectation of a quick response time that is dependent on the degradation rates of the intermediate RNA species that propagate the signals in the network. Our simple estimate shows that for circuits constructed from the attenuation system, we can expect the response time to be ∼ 5 minutes for each step in the circuit. This is in stark contrast to an analogous protein-mediated cascade, which has been shown to have a response time of ∼ 140 minutes as measured *in vivo* ^44^. In fact, because protein degradation is typically very slow, the response times of protein-mediated circuitry is often set by the rate of cell division, which is the dominant source of protein decay ^26^.

### Cascade response time can be tuned by using tandem attenuators

The success of the cascade response time measurements led us to begin the forward design of an RNA version of the single input module (SIM) ^37^. The SIM is a network motif in which a single regulatory molecule controls the expression of multiple outputs (Figure 5A). In nature, SIMs are used to regulate the genes of biosynthetic pathways and stress response systems so that they are expressed in the order at which they are needed ^2^. The SIM provides this temporal regulation by encoding different regulatory thresholds of each gene. As the master regulator increases in concentration it successively traverses these thresholds and thus activates genes at different times ^37^ (Figure 5A).

**Figure 5.**
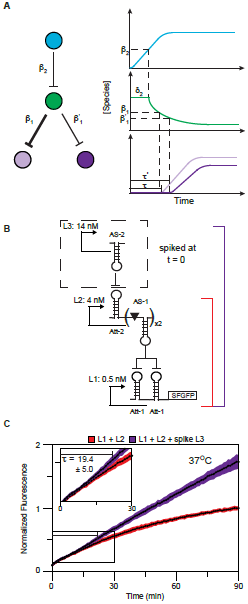
Determining the response time to tandem attenuators. (A) Schematic of a single input module (SIM). Colored circles represent nodes of the circuit, a repression cascade in which two genes (purple circles) are temporally controlled by a single species (blue circle). Temporal control is accomplished by varying thresholds of action (β1, β1’) for the intermediate cascade species (green circle). Plots are schematic response curves for the spike experiment described in Figure 4. The regulatory species (blue) is added in at t = 0. This species shuts off production of the intermediate species (green) once threshold β2 is reached at time δ2 (see Supplementary Appendix 1). The final genes are activated after time τ and τ’ once the intermediate species falls below the thresholds β1 and β1’, respectively. (B) Schematic of spike experiment. Experimental setup was analogous to Figure 4A except for L1 containing tandem pT181 attenuators (Att-Att). (C) Normalized fluorescence curves combining three separate experiments performed at 37°C with a total of 12 replicates over multiple days. L1 (0.5 nM) + L2 (4nM) reaction was setup for 20 min at which point L3 (14 nM, purple curve) or control DNA (14 nM, red curve) was spiked into the reaction and time reset to 0. Inset shows the response time of the circuit to the addition of L3 (τ = 19.4 ± 5.0 min). Shaded regions represent standard deviations calculated at each time point.

The ability to configure tandem attenuators upstream of a coding sequence provides a mechanism to adjust the threshold at which the RNA cascade responds to antisense concentrations (Figure 5B). Since tandem attenuators are more sensitive to antisense RNA concentration ^18^, we hypothesized that the response time of a cascade with tandem attenuators in L1 (Att-Att) would be slower than that of the single attenuator cascade –i.e. it would take a longer time for AS-1 to decay below the threshold required for repressing the Att-Att attenuator, thus causing a longer response time (Figure 5A). We repeated the L3 spike experiment with the Att-Att cascade at 37°C and determined the response time to be 19.4 ± 5.0 min (Figure 5C, Supporting Table S1). This response time was statistically significant from the single Att cascade at 37°C (p = 0.0303, Figure 4C) and again matched the estimation of 20 minutes based on a modified version of Equation 1 that takes into account the different threshold of the tandem Att-Att (Supporting Appendix 1). We thus showed that the Att-Att tandem attenuator could be effectively used to tune cascade response time, making it suitable for its use as a component in designing an RNA SIM.

### A theophylline responsive antisense provides a bridge to move RNA circuitry *in vivo*

While the tandem attenuator cascade provided a necessary component of the RNA SIM, we have thus far been probing circuit response time by spiking in antisense RNA via a DNA plasmid - a perturbation not possible in an *in vivo* experiment. We therefore changed L3 to encode the theophylline-responsive antisense RNA developed by Qi et al. ^19^, which has previously been shown to only attenuate transcription in the presence of theophylline *in vivo*. The response time probing experiment was adjusted so that either theophylline (2 mM in the reaction) or water (as a control) was spiked into each TX-TL reaction containing L1, L2, and aptamer-L3 (Figure 6A). The response time was then calculated by comparing with (+) and without (-) theophylline fluorescence trajectories in the same way as described above. This was done for both the single (Att) and tandem (Att-Att) versions of the cascade resulting in response times of 59.3 ± 7.3 min and 45.2 ± 11.7 min respectively (Figure 6B-C, Supporting Table S1).

**Figure 6.**
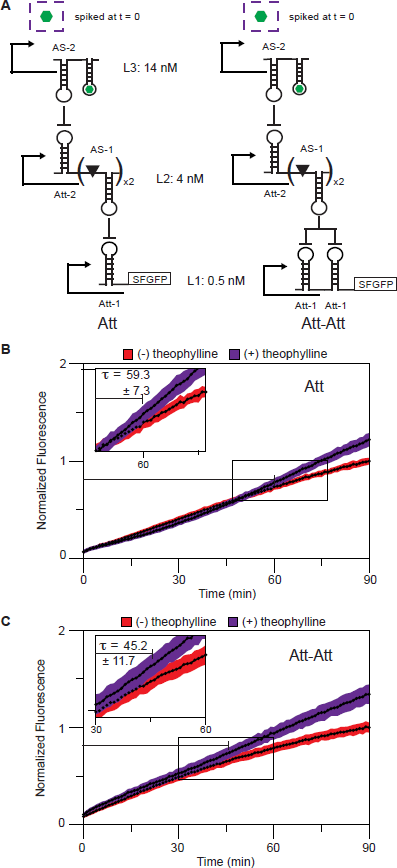
Determining the response time to a theophylline regulated cascade. (A) Schematic of experiment. L3 of the cascade in Figure 4A and 5A has been replaced with AS-2 fused to a theophylline aptamer 19. AS-2 is only active in the presence of theophylline. Theophylline was spiked into an ongoing L1 + L2 + aptamer-L3 TX-TL reaction at t = 0 (represented by dashed box). (B) Single attenuator (Att) cascade. Normalized fluorescence curves combining three separate experiments performed at 37°C with a total of 12 replicates over multiple days. L1 (Att, 0.5 nM) + L2 (4nM)+aptamer L3 (14 nM) reaction was setup for 20 min at which point theophylline (final concentration 2 mM, puple curve) or ddH2O (red curve) was spiked into the reaction and time reset to 0. Inset shows the response time of the circuit to the addition of theophylline (τ = 59.3 ± 7.3 min). (C) Tandem attenuator (Att-Att) cascade. Normalized fluorescence curves combining three separate experiments performed at 37°C with a total of 9 replicates over multiple days. L1 (Att-Att, 0.5 nM) + L2 (4nM) + aptamer L3 (14 nM) reaction was setup for 20 min at which point theophylline (final concentration 2 mM, puple curve) or ddH2O (red curve) was spiked into the reaction and time reset to 0. Inset shows the response time of the circuit to the addition of theophylline (τ = 45.2 ± 11.7 min). Shaded regions represent standard deviations calculated at each time point.

Notably, both response times are greater than what we observed in the DNA spike experiments (Figures 4,5). From Equation (1) we see that this response time is governed by the degradation rates of the two intermediate species. There are two major differences between the DNA and theophylline spike experiments that can affect these degradations rates. The first is a buildup of a pool of unbound aptamer-AS-2 in the latter case, which competes for degradation machinery slowing down overall degradation rates. The second difference is the presence of the extra aptamer hairpin on the AS-2 RNA, which could have the effect of stabilizing this RNA ^17^. These effects could result in a slower degradation rate and thus slower response time.

Interestingly, we also observe that the response time for the Att cascade was longer than that of the Att-Att cascade in the theophylline spike experiments. We attribute this to the dip in the Att fluorescence trajectory with (+) theophylline curve seen between 0-30 min (Figure 6B, Supporting Figure S6), that is not present in the Att-Att (+) trajectory (Figure 6C), causing a larger response time to be calculated with our method. In the theophylline spike experiments, we hypothesize that the presence of unbound aptamer-AS-2 competes with AS-1 for RNA degradation machinery. Because of the high concentration of aptamer-L3 used in these experiments, the aptamer-L3 RNAs would be present in a higher concentration than AS-1, and could also be more stable due to the addition of an extra hairpin ^17^. This bottleneck in RNA degradation could cause a dip in the fluorescence trajectory – the SFGFP production rate could transition from a slow to a fast phase once the excess unbound aptamer-L3 is cleared from the reaction allowing AS-1 to be degraded and the cascade to be fully activated. The presence of this dip confounds our calculation of circuit response time, leading to a larger response time estimation. From this reasoning, we would also expect to see a dip in the Att-Att (+) theophylline curve, which we do not observe (Figure 6B). It could be that this dip is more subtle, and therefore and is masked by the noise of the experiments. This may be expected because of the overall lower SFGFP production rate from the Att-Att construct 18 causing changes in this rate to be harder to observe.

Another difference that could account for the dip in the Att fluorescence trajectory is that in the theophylline experiments, the aptamer-L3 construct is present throughout the entire reaction. This means that an extra 14 nM of DNA is in the reaction using up TX-TL resources for the 20 min pre-spike duration. This resource depletion would be worse in the Att case because of higher expression of mRNA and SFGFP from the Att construct compared to the Att-Att leading to a longer than expected response time.

Despite not observing the anticipated difference in response time between the two theophylline responsive cascades, we moved forward with the RNA SIM construction and characterization *in vivo*. While TX-TL is meant to serve as a model for conditions *in vivo* ^27^, RNA degradation machinery could be faster or more abundant in cells, or other conditions different, which would allow for proper functioning of the theophylline cascades.

### An RNA SIM functions *in vivo*

Our success in demonstrating different response times using single and tandem attenuators in TX-TL led us to design an RNA SIM and characterize its function *in vivo*. To construct the SIM, we combined the single and tandem attenuator cascades in a single circuit controlling the expression of two different fluorescent proteins, red fluorescent protein (RFP) and SFGFP respectively (Figure 7A). With the single attenuator controlling RFP and the double attenuator controlling SFGFP, we may expect that SFGFP expression would turn ON approximately 5 min later than RFP expression based on the observed response time differences in our TX-TL DNA spike experiments (Figure 4C and 5B). However, because the approximate maturation times of the RFP and SFGFP proteins are 42 min ^45^ and 5 min ^43^, respectively, we actually expected to observe the SFGFP signal to turn ON first after approximately 20 minutes, and the RFP signal to turn ON later at approximately 52 minutes, or approximately 30 min after the SFGFP signal.

**Figure 7.**
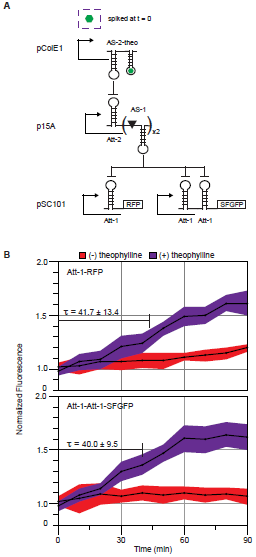
An RNA single input module (SIM) functions in vivo. (A)Schematic of the network motif. L1 contains Att-1-RFP and Att-1-Att-1-SFGFP. Expression of both proteins is controlled by AS-1 in L2, which in turn is controlled by the interaction of Att-2 with its antisense AS-2 (L3). AS-2 is a fusion with the theophylline aptamer, which is only active when the aptamer is in the bound state 19. All three plasmids were co-transformed into E. coli TG1 cells. Plasmid origins are noted by the levels. Theophylline is added to one of the split cultures once in logarithmic growth at which point time was set to zero (represented by dashed box). (B) Normalized fluorescence time courses for cultures with (+) and without (-) theophylline at 2 mM. Response time, τ, was calculated by determining the time at which the (+) and (-) curves were statistically different. τ (RFP) = 41.7 ± 13.4 min. τ (SFGFP) = 40.0 ± 9.5 min. Shaded regions represent standard deviations calculated from 12 independent transformants at each time point.

The first step in implementing the proposed RNA SIM was to develop a three-plasmid version of the cascade. To do this, we placed L1 on a low copy pSC101 backbone with kanamycin resistance, L2 on a medium copy p15A backbone with chloramphenicol resistance, and L3 on a high copy ColE1 backbone with ampicillin resistance (Supporting Figure S7A). A steady-state test of this circuit in *E. coli* TG1 cells showed 68% attenuation with L1 + L2 present, and recovery of 68% of the L1 only signal with the full cascade present (Supporting Figure S7B). Differences between these attenuation and recovery levels, and those observed in TX-TL (Figure 2F) could be due to plasmid concentrations not obeying the same DNA concentration ratios that we used in TX-TL. However, our observation of recovery upon adding L3 *in vivo* was sufficient to measure the response time of the circuit.

To construct the SIM in the three-plasmid architecture, we placed a single pT181 attenuator in front of the RFP coding sequence followed by tandem pT181 attenuators in front of the SFGFP coding sequence on the same pSC101 backbone, each under the control of its own promoter (Supporting Figure S8). This plasmid was co-transformed into *E. coli* TG1 cells along with the L2 and aptamer-L3 plasmids to complete the SIM motif. After overnight growth of replicate colonies, cultures were split in pairs and subcultured in minimal media for 4 h until logarithmic growth was reached. At this point, theophylline was added to one of the subcultures from each colony to a final concentration of 2 mM. RFP and SFGFP fluorescence was then monitored every 10 min for a total of 90 min (see Materials and Methods). RFP and SFGFP fluorescence trajectories for the with and without theophylline conditions are shown in Figure 7B. From these curves, we calculate a response time of 41.7 ± 13.4 min for Att-RFP and 40.0 ± 9.5 min for Att-Att-SFGFP.

While the Att-RFP response time is within the error of our estimate of 52 min, the Att-Att response time is outside of our estimate of 20 min (Supporting Appendix 1). Discrepancies in our estimates and measured response times could be due to three factors: i) the large standard deviations in the measurements caused by the low resolution in our time points, ii) differences due to the use of the theophylline-responsive antisense (Figure 6), iii) and fundamental differences between TX-TL reactions and *in vivo* gene expression such as the presence of other cellular factors and different cellular growth conditions. Despite these discrepancies, after accounting for fluorescent protein maturation time, we observe a clear difference in response time between the Att and Att-Att components of the RNA SIM in *E. coli*. This network motif allows for temporal control of two genes in response to a theophylline signal, and could have applications in a variety of contexts where this level of control could be useful such as optimizing metabolic pathways ^46^.

## Conclusions

In conclusion, we have demonstrated the utility of TX-TL reactions for rapidly prototyping and characterizing RNA circuitry. Each of the TX-TL experiments performed took a matter of hours to complete, and if DNA constructs are already available, several experiments can be completed with TX-TL in a single day. This is a fundamental speed up of the biological design-build-test cycle, and demonstrates the power of TX-TL as a bridge to engineering fully functioning genetic circuitry that operates *in vivo*.

In addition to establishing TX-TL reactions as a way to characterize RNA circuitry, we have also used it to directly measure circuit response times with DNA spike experiments. With these experiments, we provide the first evidence that RNA circuits can propagate signals at timescales set by their degradation rates, and showed that this leads to circuit response times on the order of 15-20 minutes. This is nearly 6 times faster than circuits constructed from stable proteins, which have response times set by the cell doubling time ^26,44^. Thus RNA circuitry shows enormous speed up compared to protein circuitry, which could become even more important in designing circuitry that needs to operate in slowly dividing cells.

We also showed how TX-TL reactions could be used to systematically prototype components for larger circuit designs. Part of our success in constructing a SIM and verifying its function *in vivo* was the result of the ability to use TX-TL to characterize its individual sub-parts and confirm that tandem attenuators could be used to tune circuit response time. This led to the construction and characterization of the first RNA SIM in *vivo*, demonstrating that RNA circuitry can be used to create temporal programs of gene expression.

Finally, we note that the initial portions of this work establishing TX-TL for characterizing RNA circuitry were performed at the inaugural Cold Spring Harbor summer course in Synthetic Biology in July-August 2013. During this intensive two week course, four of us (VS, KJS, SAK and CPF), who had little to no prior experience in performing TX-TL reactions or engineering RNA circuitry, were able to confirm the functioning of the RNA cascade and prototype a version of the RNA SIM. The rapid timescale of TX-TL experiments for characterizing genetic circuitry, and its simple experimental design ^27^ thus provides an ideal tool to teach the next generation of synthetic biologists in a cutting-edge research setting.

## Methods

### Plasmid construction and purification

A table of all the plasmids used in this study can be found in Supporting Table S4, with key sequences found in Supporting Tables S3. The pT181 attenuator and antisense plasmids, pT181 mutant attenuator and antisense plasmids, and the no-antisense control plasmid were constructs pAPA1272, pAPA1256, pAPA1273, pAPA1257, and pAPA1260, respectively, from Lucks et al. ^18^. The theophylline pT181 mutant antisense plasmid was construct pAPA1306 from Qi et al. ^19^. The second level of the cascade was modified from construct pAPA1347 from Lucks et al. ^18^. The double attenuator and SIM constructs were created using Golden Gate assembly ^47^. Plasmids were purified using a Qiagen QIAfilter Plasmid Midi Kit (Catalog number: 12243) followed by isopropanol precipitation and eluted with double distilled water.

### TX-TL extract and buffer preparation

#### Extract preparation

Cell extract and reaction buffer was prepared according to Shin and Noireaux ^36^ and Sun et al ^27^. In brief, E. coli BL21 Rosetta cells were grown to an OD600 of 1.5, pelleted via centrifugation, and washed with a buffer at pH 7.7 containing Mg-glutamate, K-glutamate, Tris and DTT. Lysis was performed via bead-beating, followed by centrifugation to remove beads and cell debris. The resulting supernatant was incubated at 37°C for 80 minutes and then centrifuged, to remove endogenous nucleic acids. The supernatant was dialyzed against a buffer at pH 8.2, containing Mg-glutamate, K-glutamate, Tris and DTT. The extract was then centrifuged, and the supernatant flash-frozen in liquid nitrogen and stored at -80°C. The cell extract for Batch 1 had a protein concentration of 37 mg/mL, and its expression was optimized via the addition of 4 mM Mg and 20 mM K. For Batch 2: 29 mg/mL protein, 4 mM Mg, and 80 mM K. Finally, Batch 3: 29 mg/mL protein, 2 mM Mg and 80 mM K.

#### Buffer preparation

The reaction buffer was prepared according to Sun et al. ^27^, and consists of an energy solution (HEPES pH 8 700 mM, ATP 21 mM, GTP 21 mM, CTP 12.6 mM, UTP 12.6 mM, tRNA 2.8 mg/ml, CoA 3.64 mM, NAD 4.62 mM, cAMP 10.5 mM, Folinic Acid 0.95 mM, Spermidine 14 mM, and 3-PGA 420 mM) and amino acids (leucine, 5 mM, all other amino acids, 6 mM).

Extract and buffer were aliquoted in separate tubes (volume appropriate for seven reactions) and stored at -80°C.

### TX-TL experiment

TX-TL buffer and extract tubes were thawed on ice for approximately 20 min. Separate reaction tubes were prepared with combinations of DNA representing a given circuit condition. Appropriate volumes of DNA, buffer, and extract were calculated using a custom spreadsheet developed by Sun et al. ^27^. Buffer and extract were mixed together and then added to each tube of DNA according to the previously published protocol ^27^. Ten μL of each TX-TL reaction mixture was transferred to a 384-well plate (Nunc 142761), covered with a plate seal (Nunc 232701), and placed on a Biotek Synergy H1 m plate reader. We note that special care is needed when pipetting to avoid air bubbles, which can interfere with fluorescence measurements. Temperature was controlled at either 29°C or 37°C. SFGFP fluorescence was measured (485 nm excitation, 520 emission) every 1-5 min depending on the experiment. Spike experiments at 29°C were paused after 25 min at which point solutions containing DNA, theophylline, or controls were added to the appropriate wells, and then placed back on the plate reader for fluorescence monitoring. Spike experiments at 37°C were paused after 20 min. In general, fluorescence trajectories were collected for 2 h, and each experiment lasting a total of 2-3 hours.

### Strains, growth media and *in vivo* gene expression

All experiments were performed in *E. coli* strain TG1. Plasmid combinations were transformed into chemically competent *E. coli* TG1 cells, plated on Difco LB+Agar plates containing 100 μg/mL carbenicillin, 34 μg/mL chloramphenicol, and 100 μg/mL kanamycin and incubated overnight at 37°C. Plates were taken out of the incubator and left at room temperature for approximately 7 h. Four colonies were picked and separately inoculated into 300 μL of LB containing carbenicillin, chloramphenicol, and kanamycin at the concentrations above in a 2 mL 96-well block (Costar 3960), and grown approximately 17 h overnight at 37°C at 1,000 rpm in a Labnet Vortemp 56 bench top shaker. Twenty μL of this overnight culture was then added to separate wells on a new block containing 930 and 980 μL (1:50 dilution) of M9 minimal media (1xM9 minimal salts, 1 mM thiamine hydrochloride, 0.4% glycerol, 0.2% casamino acids, 2 mM MgSO_4_, 0.1 mM CaCl_2_) containing the selective antibiotics and grown for 4 h at the same conditions as the overnight culture. Two wells were used for the M9 growth to represent (+) and (-) theophylline conditions for the same colony. At this point, 50 μL of a 40 mM theophylline solution was added to the wells containing 930 μL of M9. The 96-well block was placed back on the shaker. Every 10 minutes for the next 90 minutes 50 μL of the cultures with and without theophylline were removed from the block and transferred to a 96-well plate (Costar 3631) containing 50 μL of phosphate buffered saline (PBS). SFGFP fluorescence (485 nm excitation, 520 nm emission), mRFP fluorescence (560 nm excitation, 630 nm emission), and optical density (OD, 600 nm) were then measured at each time point using a Biotek Synergy H1 m plate reader.

### Response Time Calculation

#### TX-TL experiments

To quantify the circuit response time, we calculated τ using data from multiple replicates in three individual experiments by first normalizing trajectories to the endpoint fluorescence of the L1+L2 condition to account for variation in fluorescence output between experiments (Supporting Figure S5). Normalized fluorescence distributions from all replicates, between each condition, were compared using Welch’s t-test at each time point to determine the time at which the L1+L2 and L1+L2+L3 data sets were statistically different from each other. The difference in average normalized fluorescence at this point was used to set a threshold, that was then used in each individual data set to determine the time at which each spiked trajectory differed from the average of the L1+L2 curves of that experiment (Supporting Figure S5). These times were then used to calculate reported τ with error.

#### In vivo experiment

To quantify the circuit response time of the SIM, we calculated τ using data from multiple replicates in three individual experiments by first normalizing trajectories to the average of the t = 0 fluorescence of each individual colony’s with and without theophylline condition. Normalized fluorescence distributions from all replicates, between each condition, were compared using Welch’s t-test at each time point to determine the time at which the with and without theophylline data sets were statistically different from each other. The difference in average normalized fluorescence at this point was used to set a threshold that was then used in each individual data set to determine the time at which each spiked trajectory differed from its corresponding no-theophylline trajectory. These times were then used to calculate reported τ with error.

## Supporting Information

Supporting Information Available: Supporting Tables 1-4, Supporting Figures 1-8, Supporting Methods and Supporting Appendix 1. This material is available free of charge via the Internet at http://pubs.acs.org.

### Abbreviations

Transcription-translation (TX-TL), single input module (SIM), super folder green fluorescent protein (SFGFP), red fluorescent protein (RFP), ribonuclease III (RNAseIII)

### Author Contributions

MKT, JC, CAH, ZZS, VS, KJS, SAK and CPF performed the experiments. MKT, JC, CAH, and JBL designed the experiments and wrote the manuscript. VA, KJS, SAK, CPF, ZZS, VN, and RM designed the experiments and edited the manuscript.

### Funding Sources

This material is based upon work supported by the National Science Foundation Graduate Research Fellowship Program [Grant No. DGE-1144153 to M.K.T.]. Defense Advanced Research Projects Agency Young Faculty Award (DARPA YFA) [N66001-12-1-4254 to J.B.L.]. Office of Naval Research Young Investigators Program Award (ONR YIP) [N00014-13-1-0531 to J. B. L.]. Defense Advanced Research Projects Agency (DARPA/MTO) Living Foundries program [HR0011-12-C-0065 to ZZS and CAH]. The CSHL Synthetic Biology course was funded by the Howard Hughes Medical Institute and the Office of Naval Research. J.B.L. is an Alfred P. Sloan Research Fellow.

## Acknowledgement

We thank Cold Spring Harbor Laboratory for hosting the inaugural Synthetic Biology summer course where portions of this work was performed. In addition, we thank the other 2013 CSHL Synthetic Biology instructors Jeff Tabor, David Savage and Karmella Haynes for their support, and the helpful comments from all of the students in the course.

## References

(1) Purnick, P. E. M., and Weiss, R. (2009) The second wave of synthetic biology: from modules to systems. Nat. Rev. Mol. Cell Biol. 10, 410–422.

(2) Alon, U. (2007) An Introduction to Systems Biology: Design Principles of Biological Circuits. Chapman and Hall/CRC, Boca Raton, Fl.

(3) Gardner, T. S., Cantor, C. R., and Collins, J. J. (2000) Construction of a genetic toggle switch in Escherichia coli. Nature 403, 339–342.

(4) Elowitz, M. B., and Leibler, S. (2000) A synthetic oscillatory network of transcriptional regulators. Nature 403, 335–338.

(5) Stricker, J., Cookson, S., Bennett, M. R., Mather, W. H., Tsimring, L. S., and Hasty, J. (2008) A fast, robust and tunable synthetic gene oscillator. Nature 456, 516–519.

(6) Tigges, M., Marquez-Lago, T. T., Stelling, J., and Fussenegger, M. (2009) A tunable synthetic mammalian oscillator. Nature 457, 309–312.

(7) Moon, T. S., Lou, C., Tamsir, A., Stanton, B. C., and Voigt, C. A. (2012) Genetic programs constructed from layered logic gates in single cells. Nature 491, 249–253.

(8) Ausländer, S., Ausländer, D., Müller, M., Wieland, M., and Fussenegger, M. (2012) Programmable single-cell mammalian biocomputers. Nature 487, 123–127.

(9) Tabor, J. J., Salis, H. M., Simpson, Z. B., Chevalier, A. A., Levskaya, A., Marcotte, E. M., Voigt, C. A., and Ellington, A. D. (2009) A Synthetic Genetic Edge Detection Program. Cell 137, 1272–1281.

(10) Basu, S., Gerchman, Y., Collins, C. H., Arnold, F. H., and Weiss, R. (2005) A synthetic multicellular system for programmed pattern formation. Nature 434, 1130–1134.

(11) Chappell, J., Takahashi, M. K., Meyer, S., Loughrey, D., Watters, K. E., and Lucks, J. (2013) The centrality of RNA for engineering gene expression. Biotechnology Journal doi: 10.1002/biot.201300018.

(12) Buskirk, A. R., Kehayova, P. D., Landrigan, A., and Liu, D. R. (2003) In vivo evolution of an RNA-based transcriptional activator. Chem Biol 10, 533–540.

(13) Isaacs, F. J., Dwyer, D. J., Ding, C., Pervouchine, D. D., Cantor, C. R., and Collins, J. J. (2004) Engineered riboregulators enable post-transcriptional control of gene expression. Nat Biotechnol 22, 841–847.

(14) Bayer, T. S., and Smolke, C. D. (2005) Programmable ligand-controlled riboregulators of eukaryotic gene expression. Nat Biotechnol 23, 337–343.

(15) Win, M. N., and Smolke, C. D. (2007) A modular and extensible RNA-based gene-regulatory platform for engineering cellular function. Proc Natl Acad Sci USA 104, 14283–14288.

(16) Rinaudo, K., Bleris, L., Maddamsetti, R., Subramanian, S., Weiss, R., and Benenson, Y. (2007) A universal RNAi-based logic evaluator that operates in mammalian cells. Nat Biotechnol 25, 795–801.

(17) Carrier, T. A., and Keasling, J. D. (1999) Library of synthetic 5’ secondary structures to manipulate mRNA stability in Escherichia coli. Biotechnol. Prog. 15, 58–64.

(18) Lucks, J. B., Qi, L., Mutalik, V. K., Wang, D., and Arkin, A. P. (2011) Versatile RNA-sensing transcriptional regulators for engineering genetic networks. Proc Natl Acad Sci USA 108, 8617–8622.

(19) Qi, L., Lucks, J. B., Liu, C. C., Mutalik, V. K., and Arkin, A. P. (2012) Engineering naturally occurring trans-acting non-coding RNAs to sense molecular signals. Nucleic acids research 40, 5775–5786.

(20) Lucks, J. B., Mortimer, S. A., Trapnell, C., Luo, S., Aviran, S., Schroth, G. P., Pachter, L., Doudna, J. A., and Arkin, A. P. (2011) Multiplexed RNA structure characterization with selective 2’-hydroxyl acylation analyzed by primer extension sequencing (SHAPE-Seq) 108, 11063–11068.

(21) Mutalik, V. K., Qi, L., Guimaraes, J. C., Lucks, J. B., and Arkin, A. P. (2012) Rationally designed families of orthogonal RNA regulators of translation. Nat Chem Biol 8, 447–454.

(22) Liu, C. C., Qi, L., Lucks, J. B., Segall-Shapiro, T. H., Wang, D., Mutalik, V. K., and Arkin, A. P. (2012) An adaptor from translational to transcriptional control enables predictable assembly of complex regulation. Nat. Methods 9, 1088–1094.

(23) Takahashi, M. K., and Lucks, J. B. (2013) A modular strategy for engineering orthogonal chimeric RNA transcription regulators. Nucleic acids research 41, 7577–7588.

(24) Xie, Z., Wroblewska, L., Prochazka, L., Weiss, R., and Benenson, Y. (2011) Multi-input RNAi-based logic circuit for identification of specific cancer cells. Science 333, 1307–1311.

(25) Qi, L. S., Larson, M. H., Gilbert, L. A., Doudna, J. A., Weissman, J. S., Arkin, A. P., and Lim, W. A. (2013) Repurposing CRISPR as an RNA-Guided Platform for Sequence-Specific Control of Gene Expression. Cell 152, 1173–1183.

(26) Rosenfeld, N., and Alon, U. (2003) Response delays and the structure of transcription networks. J Mol Biol 329, 645–654.

(27) Sun, Z. Z., Hayes, C. A., Shin, J., Caschera, F., Murray, R. M., and Noireaux, V. (2013) Protocols for Implementing an Escherichia coli Based TX-TL Cell-Free Expression System for Synthetic Biology. Journal of Visualized Experiments, 79, e50762.

(28) Shin, J., and Noireaux, V. (2012) An E. coliCell-Free Expression Toolbox: Application to Synthetic Gene Circuits and Artificial Cells. ACS Synth. Biol. 1, 29–41.

(29) Karig, D. K., Iyer, S., Simpson, M. L., and Doktycz, M. J. (2012) Expression optimization and synthetic gene networks in cell-free systems. Nucleic acids research 40, 3763–3774.

(30) Niederholtmeyer, H., Stepanova, V., and Maerkl, S. J. (2013) Implementation of cell-free biological networks at steady state. Proc Natl Acad Sci USA 110, 15985–15990.

(31) Hodgman, C. E., and Jewett, M. C. (2012) Cell-free synthetic biology: Thinking outside the cell. Metabolic Engineering 14, 261–269.

(32) Chappell, J., Jensen, K., and Freemont, P. S. (2013) Validation of an entirely in vitro approach for rapid prototyping of DNA regulatory elements for synthetic biology. Nucleic acids research 41, 3471–3481.

(33) Noireaux, V., Bar-Ziv, R., and Libchaber, A. (2003) Principles of cell-free genetic circuit assembly. Proc Natl Acad Sci USA 100, 12672–12677.

(34) Shimizu, Y., Inoue, A., Tomari, Y., Suzuki, T., Yokogawa, T., Nishikawa, K., and Ueda, T. (2001) Cell-free translation reconstituted with purified components : Abstract : Nature Biotechnology. Nat Biotechnol 19, 751–755.

(35) Sun, Z. Z., Yeung, E., Hayes, C. A., Noireaux, V., and Murray, R. M. (2013) Linear DNA for Rapid Prototyping of Synthetic Biological Circuits in an Escherichia coli Based TX-TL Cell-Free System. ACS Synth. Biol. doi: 10.1021/sb400131a.

(36) Shin, J., and Noireaux, V. (2010) Efficient cell-free expression with the endogenous E. Coli RNA polymerase and sigma factor 70. J Biol Eng 4, 8.

(37) Shen-Orr, S. S., Milo, R., Mangan, S., and Alon, U. (2002) Network motifs in the transcriptional regulation network of Escherichia coli. Nat Genet 31, 64–68.

(38) Novick, R. P., Iordanescu, S., Projan, S. J., Kornblum, J., and Edelman, I. (1989) pT181 plasmid replication is regulated by a countertranscript-driven transcriptional attenuator. Cell 59, 395–404.

(39) Brantl, S., and Wagner, E. G. (2000) Antisense RNA-mediated transcriptional attenuation: an in vitro study of plasmid pT181. Mol Microbiol 35, 1469–1482.

(40) Kolb, F. A., Engdahl, H. M., Slagter-Jäger, J. G., Ehresmann, B., Ehresmann, C., Westhof, E., Wagner, E. G., and Romby, P. (2000) Progression of a loop-loop complex to a four-way junction is crucial for the activity of a regulatory antisense RNA. EMBO J 19, 5905–5915.

(41) Nicholson, A. W. (1999) Function, mechanism and regulation of bacterial ribonucleases. FEMS Microbiology Reviews 23, 371–390.

(42) Brantl, S., and Wagner, E. G. H. (2002) An Antisense RNA-Mediated Transcriptional Attenuation Mechanism Functions in Escherichia coli. Journal of Bacteriology 184, 2740–2747.

(43) Pédelacq, J.-D., Cabantous, S., Tran, T., Terwilliger, T. C., and Waldo, G. S. (2006) Engineering and characterization of a superfolder green fluorescent protein. Nat Biotechnol 24, 79–88.

(44) Hooshangi, S., Thiberge, S., and Weiss, R. (2005) Ultrasensitivity and noise propagation in a synthetic transcriptional cascade. Proc Natl Acad Sci USA 102, 3581–3586.

(45) Campbell, R. E., Tour, O., Palmer, A. E., Steinbach, P. A., Baird, G. S., Zacharias, D. A., and Tsien, R. Y. (2002) A monomeric red fluorescent protein. Proc Natl Acad Sci USA 99, 7877–7882.

(46) Zaslaver, A., Mayo, A. E., Rosenberg, R., Bashkin, P., Sberro, H., Tsalyuk, M., Surette, M. G., and Alon, U. (2004) Just-in-time transcription program in metabolic pathways. Nat Genet 36, 486–491.

(47) Engler, C., Kandzia, R., and Marillonnet, S. (2008) A one pot, one step, precision cloning method with high throughput capability. PLoS ONE 3, e3647.

